# Extensive genetic differentiation between homomorphic sex chromosomes in the mosquito vector, *Aedes aegypti*

**DOI:** 10.1101/060061

**Authors:** Albin Fontaine, Igor Filipović, Thanyalak Fansiri, Ary A. Hoffmann, Changde Cheng, Mark Kirkpatrick, Gordana Rašić, Louis Lambrechts

## Abstract

Mechanisms and evolutionary dynamics of sex-determination systems are of particular interest in insect vectors of human pathogens like mosquitoes because novel control strategies aim to convert pathogen-transmitting females into non-biting males, or rely on accurate sexing for the release of sterile males. In *Aedes aegypti,* the main vector of dengue and Zika viruses, sex determination is governed by a dominant male-determining locus, previously thought to reside within a small, non-recombining, sex-determining region (SDR) of an otherwise homomorphic sex chromosome. Here, we provide evidence that sex chromosomes in *Ae. aegypti* are genetically differentiated between males and females over a region much larger than the SDR. Our linkage mapping intercrosses failed to detect recombination between X and Y chromosomes over a 123-Mbp region (40% of their physical length) containing the SDR. This region of reduced male recombination overlapped with a smaller 63-Mbp region (20% of the physical length of the sex chromosomes) displaying high male-female genetic differentiation in unrelated wild population from Brazil and Australia and in a reference laboratory strain originating from Africa. In addition, the sex-differentiated genomic region was associated with a significant excess of male-to-female heterozygosity and contained a small cluster of loci consistent with Y-specific null alleles. We demonstrate that genetic differentiation between sex chromosomes is sufficient to assign individuals to their correct sex with high accuracy. We also show how data on allele frequency differences between sexes can be used to estimate linkage disequilibrium between loci and the sex-determining locus. Our discovery of large-scale genetic differentiation between sex chromosomes in *Ae. aegypti* lays a new foundation for mapping and population genomic studies, as well as for mosquito control strategies targeting the sex-determination pathway.

## Introduction

Understanding the underlying mechanisms and evolutionary dynamics of sex determination in mosquitoes is of particular interest as new strategies for controlling mosquito-borne diseases aim to convert pathogen-transmitting females into non-biting males (Hall, et al. 2015), or rely on accurate sexing for the release of sterile males (Eckermann, et al. 2014; Gilles, et al. 2014). Sex determination in mosquitoes and other dipterans is under the control of a gene regulation cascade that relies on alternative splicing of genes expressed in both males and females (Salz 2011). There is a great variation across dipteran species, and even between populations of the same species, in how this cascade is initiated (Bopp, et al. 2014). The master switch at the top of the cascade in drosophilids is the number of X chromosomes, whereas in tephritids, houseflies and mosquitoes it is a dominant male-determining factor (Kaiser and Bachtrog 2010; Vicoso and Bachtrog 2015).

In *Aedes* and *Culex* mosquitoes, the male-determining locus is located on a morphologically undifferentiated (homomorphic) sex chromosome, which is considered the ancestral state of mosquito sex chromosomes (Toups and Hahn 2010). In contrast, the malarial mosquitoes (Anophelinae) have acquired fully morphologically differentiated (heteromorphic) X and Y chromosomes (Toups and Hahn 2010). Why heteromorphy of sex chromosomes evolved in some mosquito lineages but not others remains unclear. Evolutionary models suggest progression of autosomes into heteromorphic sex chromosomes after the acquisition of a sex-determining locus (Charlesworth 1996; Charlesworth and Charlesworth 2000). According to these models, the selective advantage of linkage between sex-determining genes and sexually antagonistic genes promotes initial suppression of recombination between homologous chromosomes, followed by expansion of the non-recombining region (Rice 1987). An evolving pair of neo-sex chromosomes further differentiates through changes in gene content, gene decay and epigenetic modifications (Bachtrog 2013). Yet, recent analyses of fly genomes revealed a striking diversity of evolutionary trajectories where sex chromosomes have been gained, lost, replaced and rearranged multiple times over the dipteran evolutionary history (Kaiser and Bachtrog 2010; Vicoso and Bachtrog 2015).

*Aedes aegypti*, the main vector of dengue, Zika, yellow fever and chikungunya viruses worldwide, has homomorphic sex chromosomes like other Culicinae (Toups and Hahn 2010). Genetic evidence suggests that the male-determining locus resides in a non-recombining, sex-determining region (SDR) of chromosome 1 (Toups and Hahn 2010). Motara and Rai (Motara and Rai 1978) proposed a nomenclature to define the copy of chromosome 1 with the M locus as the M chromosome, and the copy without the M locus as the m chromosome. Thereafter, we follow the standard terminology and refer to the m and M chromosomes as X and Y chromosomes, respectively. Motara and Rai also noticed some cytological differences consistent with differentiation of the SDR between the X and Y chromosomes of *Ae. aegypti* (Motara and Rai 1978). However, fine details of chromosomal features have been elusive due to problems in producing high-quality, easily spreadable polytene chromosomes in *Ae. aegypti* (Timoshevskiy, et al. 2013).

Availability of a reference genome sequence (Nene, et al. 2007) and affordable high-throughput sequencing technologies have opened new avenues to characterize genomic features in *Ae. aegypti*. Produced nearly 10 years ago, the *Ae. aegypti* reference genome sequence encompasses 1.39 billion base pairs (Gbp) fragmented in over 4,700 scaffolds (Nene, et al. 2007) that were recently assembled into end-to-end chromosomes by chromosome conformation capture (Dudchenko et al. 2017). Prior to this chromosome-wide assembly, linkage mapping using restriction site-associated DNA sequencing (RADseq) and physical mapping by fluorescent *in situ* hybridization (FISH) with metacentric chromosome preparations were used to produce partial assemblies, with up to two thirds of the genome sequence assigned to distinct chromosomes (Juneja, et al. 2014; Timoshevskiy, et al. 2013). Recent comparative genomic analyses suggested a particularly dynamic evolution of sex chromosomes that contain synteny blocks with the X and 2R chromosome arms of *Anopheles gambiae* (Timoshevskiy, et al. 2014).

The homomorphy of *Ae. aegypti* sex chromosomes was also inferred from genome-wide sequencing coverage differences between males and females (Vicoso and Bachtrog 2015). Because females have two copies of the X chromosome and males have only one, X-specific scaffolds are expected to display about half the depth of sequencing coverage in males compared to females. Vicoso and Bachtrog (2015) did not find a significant difference in depth of coverage between *Ae. aegypti* males and females when analyzing paired-end Illumina reads from whole-genome sequencing (WGS) libraries. This indicated that the Y-chromosome sequences are not sufficiently divergent from the X-chromosome sequences to preclude their successful alignment to the reference scaffolds, thus supporting the existence of undifferentiated sex chromosomes in *Ae. aegypti.* Hall and colleagues (2014) used a similar approach called the chromosome quotient method to identify the male-determining gene(s) within the SDR, but failed to do so when using the current *Ae. aegypti* genome sequence. Instead, they identified the male-determining gene *Nix* (Hall, et al. 2015) and male-biased sequences primarily found in the male genome, such as the gene *myo-sex* (Hall, et al. 2014), from transcriptomic data, expressed sequence tags and unassembled bacterial artificial chromosomes.

The overall conclusions from the previous studies are that: (*i*) the SDR in *Ae. aegypti* occupies a small region that maps to band 1q21 of chromosome 1, (*ii*) regions near the SDR (including *myo-sex*) show low levels of recombination with the X chromosome (Hall, et al. 2014), and (*iii*) most of chromosome 1 recombines in an autosome-like fashion thereby maintaining the overall homomorphy. Also, the current genome sequence is considered largely uninformative when looking for sequences primarily found in the male genome (Hall, et al. 2015; Hall, et al. 2014).

Here, we provide compelling genetic evidence that, despite the apparent homomorphy, sex chromosomes in *Ae. aegypti* are genetically differentiated over a region much larger than the non-recombining SDR. Our linkage mapping experiments failed to detect recombination in F_1_ males over a 123-million-base-pair (Mbp) region of chromosome 1, spanning from position 87 Mbp to position 210 Mbp and representing about 40% of its physical length. Analyses of genome-wide variation in the unrelated laboratory strain (Liverpool) used to generate the current reference genome sequence, as well as in wild *Ae. aegypti* populations revealed substantial male-female genetic differentiation in a smaller 63-Mbp region spanning from position 148 Mbp to position 211 Mbp and representing about 20% of the physical length of the sex chromosomes. A small cluster of loci located inside this region displayed genotypic patterns consistent with Y-specific null alleles. We further show that genetic differentiation between sex chromosomes is sufficient to accurately assign individuals to their phenotypic sex. We also demonstrate that allele frequency differences between males and females can be used to estimate linkage disequilibrium (LD) with the SDR. Our results lay a new foundation for the mapping and population genomic studies in *Ae. aegypti*, and for the control strategies that rely on accurate sexing and sex reversal in this important mosquito vector.

## Materials and Methods

### Mosquito Samples for Laboratory Crosses

Two independent laboratory crosses were carried out with wild-type *Ae. aegypti* mosquitoes originally collected in February 2011 from Kamphaeng Phet, Thailand. Cross #1 was an F_2_ intercross created with a single virgin male from one isofemale line and a virgin female from another isofemale line. Both isofemale lines were derived from wild *Ae. aegypti* founders from Thailand collected as eggs using ovitraps as previously described (Fansiri, et al. 2013). Prior to the cross, the lines were maintained in the laboratory by mass sib-mating and collective oviposition until the 19^th^ generation. This was done to increase homozygosity and maximize the number of informative markers for our linkage mapping design. A total of 22 males and 22 females from the Cross #1 F_2_ progeny were used to generate a linkage map and subsequently map the sex-determining locus. Cross #2 was an F_2_ intercross between a pair of field-collected mosquito founders from Thailand (Fansiri, et al. 2013). Adults were maintained in an insectary under controlled conditions (28± 1°C, 75±5% relative humidity and 12:12 hour light-dark cycle). The male and female of each mating pair were chosen from different collection sites to avoid sampling siblings (Apostol, et al. 1994; Rašić, et al. 2016; Rasic, et al. 2014). Egg batches from the same F_0_ female were merged to obtain a pool of F_1_ eggs and F_2_ progeny was produced by mass sib-mating and collective oviposition of the F_1_ offspring (Supplementary File 1E). A total of 197 females of the Cross #2 F_2_ progeny were used to generate a linkage map.

### Field Samples for Population Genomic Analyses

Field-caught *Ae. aegypti* samples from Rio de Janeiro, Brazil and Queensland, Australia were analyzed. Samples of 62 adult mosquitoes from Australia were caught using Biogents sentinel traps set up in Gordonvale (17 females and 17 males) and Townsville (14 females and 14 males), Queensland in January 2014 (Rašić, et al. 2016). Adults were identified as males or females based on the sexually dimorphic antennae and external genitalia structure (Becker 2003). Mosquitoes from Rio de Janeiro, Brazil were collected from ovitraps in November-December 2011 (Rašić, et al. 2015). Larvae were reared until the third instar in an insectary under controlled conditions (25± 1°C, 80±10% relative humidity and 12:12 hour light-dark cycle). Only one individual per ovitrap was retained to avoid analyzing siblings. Sex of each individual was determined based on the presence or absence of the highly male-biased sequence *myo-sex* (Hall, et al. 2014) and confirmed with two additional male-specific sequences that were identified in this study (Supplementary File 1A,C). The final dataset from Brazil consisted of 66 mosquitoes (32 females and 34 males).

### DNA Extraction

Mosquito genomic DNA was extracted using the NucleoSpin 96 Tissue Core Kit (Macherey-Nagel, Düren, Germany). To obtain a sufficient amount of DNA for the parental males from the laboratory crossings, whole-genome amplification was performed by multiple displacement amplification using the Repli-g Mini kit (Qiagen, Hilden, Germany). All DNA concentrations were measured with Qubit fluorometer and Quant-iT dsDNA Assay kit (Life technologies, Paisley, UK).

### Double-digest RADseq Library Generation

An adaptation of the original double-digest restriction-site associated DNA sequencing (ddRADseq) protocol (Peterson, et al. 2012) was used as previously described (Rasic, et al. 2014). Briefly, 500 ng of genomic DNA from each mosquito was used for the mapping samples, and 100 ng for the field-collected samples. DNA was digested in with *NlaIII* and *MluCI* restriction enzymes (New England Biolabs, Herts, UK), for 3 hours at 37°C. Cleaned digestions were ligated to the modified Illumina P1 and P2 adapters with overhangs complementary to *NlaIII* and *MluCI* cutting sites, respectively. Each mosquito was uniquely labeled with a combination of P1 and P2 barcodes. Ligation reactions were incubated at 16°C overnight and heat inactivated. Adapter ligated DNA fragments from all individuals were then pooled and cleaned with 1.5× bead solution. Size selection of fragments between 350–440 base pairs (bp) for the laboratory crosses or 300-450 bp for the field populations was performed using a Pippin-Prep 2% gel cassette (Sage Sciences, Beverly, MA, USA). Finally, 1 μl of the size-selected DNA was used as a template in a 10-μl PCR reaction. To reduce PCR duplicates bias, 8 PCR reactions were run in parallel, pooled, and cleaned to make the final library. Final libraries were quantified by quantitative PCR using the QPCR NGS Library Quantification Kit (Agilent technologies, Palo Alto, CA, USA). For the mapping crosses, libraries spiked with 15% PhiX were sequenced in paired-end on an Illumina NextSeq 500 platform using a 150-cycle chemistry (Illumina, San Diego, CA, USA) (accession numbers pending). For the field populations, four ddRADseq libraries spiked with 10% PhiX were sequenced in paired-end on an Illumina HiSeq 2000 platform with a 100-cycle chemistry (NCBI SRA accession numbers SRX1970106-108, SRX2248021).

### Bioinformatics Processing and Genotype Calling

A previously developed bash script pipeline (Rasic, et al. 2014) was used to process raw sequence reads. Briefly, the DDemux program was used for demultiplexing fastq files according to the P1 and P2 barcodes combinations. Sequences were filtered with FASTX-Toolkit, discarding the reads with Phred scores < 25. Reads were trimmed to 90 bp (HiSeq platform) and 140 bp (NextSeq platform) on both P1 and P2 ends. Reads were then aligned to the reference *Ae. aegypti* genome (AaegL1, February 2013) (Nene, et al. 2007) using Bowtie version 0.12.7 (Langmead, et al. 2009). Parameters for the ungapped alignment included a maximum of three mismatches allowed in the seed, suppression of alignments if more than one reportable alignment exists, and a “try-hard” option to find valid alignments. Aligned Bowtie output files were imported into the Stacks pipeline (Catchen, et al. 2013; Catchen, et al. 2011).

A catalogue of RAD loci used for single nucleotide polymorphism (SNP) discovery was created using the ref_map.pl pipeline in Stacks version 1.19 (Catchen, et al. 2013; Catchen, et al. 2011). A RAD locus was generated with a minimum of 5 reads. For the mapping crosses, the “genotypes” module was used to export F_2_ mosquito genotypes for all markers homozygous for alternative alleles in the F_0_ parents (i.e., homozygous AA in the F_0_ male and homozygous BB in the F_0_ female) with a sequencing depth ≥12× in ≥60% of the mapping population, to minimize the risk of false homozygous calls.

AaegL1 genomic coordinates were translated into the recently published chromosome-wide AaegL4 assembly coordinates (Dudchenko et al. 2017). The AaegL1 assembly was blasted using blastn v2.6.0 against the AaegL4 assembly with default parameters, except a word size of 1,000 bp and a percentage identity of 100% between query and subject sequences. The genomic coordinates of each marker were translated based on the blast output file using an in-house awk script that accounted for potential fragment inversions between the two assemblies.

### Linkage Map Construction

A comprehensive linkage map based on recombination fractions among RAD markers in the F_2_ generation was constructed using the R package OneMap v2.0-3 (Margarido, et al. 2007). Marker positions on the chromosome-wide AaegL4 assembly facilitated the assignment of markers to a linkage group and their respective ordering. Following the selection of markers for which the parents were homozygous for alternative alleles, in Cross #1 every independent marker was expected to segregate in the F_2_ mapping population at a frequency of 25% for homozygous genotypes (AA and BB) and of 50% for heterozygous genotypes (AB) when considering both males and females together. A 𝒳^2^ test was used to filter out markers based on deviations of the observed genotype frequencies in the F_2_ progeny from the Mendelian segregation ratios expected for autosomal loci. Fully sex-linked markers are expected to segregate in F_2_ females with equal frequency (50%) of AB and BB genotypes, because the F_0_ paternal AA genotype only occurs in F_2_ males (Supplementary File 1D). Reciprocally, fully sex-linked markers in F_2_ males are expected to lack F_0_ maternal BB genotypes.

Cross #2 could only be analyzed as a classical F_2_ intercross design for autosomal linkage groups because only females were genotyped. Markers located on autosomes were filtered out based on deviations from expected Mendelian segregation ratios as described above. Markers on chromosome 1 were included if they had heterozygous genotype (AB) frequencies inside the]40% – 60%[range and F_0_ maternal genotype (BB) frequencies inside the]5% – 65%[range. These arbitrary limits for initial marker selection were largely permissive for pseudo-autosomal markers segregating according to theoretical proportions (0-25% AA: 50% AB: 25% – 50% BB). To our knowledge, there is no linkage analysis method readily available to deal with such sex-specific deviations in genotype segregation ratios. Linkage analysis in Cross #2 was therefore restricted to chromosomes 2 and 3.

Recombination fractions between all pairs of selected markers were estimated using the rf.2pts function with default parameters. Because sex-specific recombination rates cannot be estimated with an F_2_ cross design, a sex-averaged recombination rate was estimated. Markers linked with a minimum LOD score of 13 and 25 for Cross #1 and Cross #2, respectively, were assigned to a same linkage group and unlinked markers were removed from further analysis. Linkage groups were assigned to the three distinct *Ae. aegypti* chromosomes based on the physical coordinates of the chromosome-wide AaegL4 assembly.

Recombination fractions were converted into genetic distances in centiMorgans (cM) using the Kosambi mapping function (Kosambi 1943). Linkage maps were exported in the R/qtl environment (Broman, et al. 2003) where they were corrected for tight double crossing-overs with the calc.errorlod function based on a LOD cutoff threshold of 2 and 1.4 for Cross #1 and Cross #2, respectively. Sex QTL detection was performed with the scanone function using a binary trait model and the EM algorithm.

### Population Genomic Analyses

Brelsford and colleagues recently demonstrated how ddRADseq can be used to identify homomorphic sex chromosomes from wild-caught adults in non-model animals (Brelsford, et al. 2016). Because males and females share the X chromosome, a maximum allele difference of 0.5 between males and females is expected when different alleles are fixed on X and Y chromosomes. In such a case, excess heterozygosity should also be observed in males when compared to females (Brelsford, et al. 2016). We used this approach to assess the extent to which chromosome 1 in unrelated wild *Ae. aegypti* populations shows enrichment for sex-differentiated markers when compared to the other two chromosomes.

RAD tags were selected that were (*i*) shared between ≥ 75% of all individuals (males and females combined) in each *Ae. aegypti* population, and (*ii*) had SNPs with a minor allele frequency ≥ 10%. Allelic difference between females and males from a population was estimated as the Weir and Cockerham *F*_ST_ statistic (Weir and Cockerham 1984). *F*_ST_ reaches a value of 0.5 for fully sex-linked markers that have alternatively fixed alleles on X and Y chromosomes. Genepop (Rousset 2008) was used to estimate *F*_ST_ and the frequency of heterozygotes (*H*) for each marker (Supplementary File 2A).

To assess if X and Y chromosomes are sufficiently differentiated to predict phenotypic sex in *Ae. aegypti*, a multivariate clustering method called discriminant analysis of principal components (DAPC) (Jombart, et al. 2010) was used in the R package “adegenet” (Jombart and Ahmed 2011). DAPC was performed separately for each population and chromosome. A discriminant function was constructed for each population to distinguish males from females, using only five retained PCs in order to avoid model over fitting (Jombart and Collins 2015). Given that DAPC finds linear combinations of allele frequencies (the discriminant functions) which best separate the clusters, sex-linked markers can be identified as those with the highest allelic contribution to the discrimination of males and females.

In addition to the wild-caught *Ae. aegypti*, genome-wide differentiation patterns between males and females from the Liverpool strain were also analyzed. This inbred line originates from West Africa and has been maintained in the laboratory since 1936 (https://www.vectorbase.org/organisms/aedes-aegypti/liverpool). The Liverpool strain was used to generate the reference genome sequence of *Ae. aegypti* (Nene, et al. 2007) as well as several partial assemblies, and remains the most commonly used material in various laboratory studies of *Ae. aegypti*. This analysis used the WGS dataset generated by Hall and colleagues (2014). Briefly, they isolated genomic DNA separately from 10 males and 6 virgin females from the Liverpool strain. A pooled WGS library for each sex was sequenced in paired-end on an Illumina HiSeq 1000 platform using a 100-cycle chemistry (NCBI SRA accession number SRP023515). Raw reads were processed and those with a quality score > 25 were aligned to the reference genome using the bash script pipeline and Bowtie parameters described above. Uniquely aligned reads were further processed and analyzed using the Popoolation2 pipeline (Kofler, et al. 2011) to estimate allele frequencies from a pooled sequencing experiment. The Weir and Cockerham *F*_ST_ statistic (Weir and Cockerham 1984) between males and females was calculated for each SNP with a depth of coverage between 50 and 200 reads.

## Results

We first calculated linkage map positions of 363 ddRADseq markers that were unambiguously ordered according to their physical position on the three linkage groups using 22 F_2_ males and 22 F_2_ females from Cross #1. The three linkage groups contained 122, 49 and 192 markers, covering 128.8, 947.6 and 232.9 cM for chromosomes 1, 2 and 3 respectively. The average spacing between markers for chromosomes 1, 2 and 3 was 1.1, 19.7 and 1.2 cM, respectively. A second linkage map spanning 129.7 cM was generated using 197 F_2_ females from Cross #2. Only female mosquitoes were genotyped in this cross because it was originally designed to map quantitative trait loci (QTL) underlying dengue vector competence, a female-specific trait. Owing to the lack of genotyped males in the F_2_ progeny and the sex-specific genotype segregation patterns (described below), it was not possible to obtain a linkage map for chromosome 1. The Cross #2 linkage map contained 61 and 77 markers with an average spacing between markers of 0.8 and 1.1 cM for chromosome 2 and 3, respectively (Supplementary Table 1).

Across both of our linkage maps generated using the ddRADseq markers, a total of 372 unique supercontigs were assigned to the three *Ae. aegypti* chromosomes (Supplementary File 2B), representing 39.6% of the base pairs from the current reference genome sequence. Linkage group assignments of supercontigs were generally in agreement with a previously published chromosome map (Juneja, et al. 2014; Timoshevskiy, et al. 2013) (Figure 1). Only 10 supercontigs (2.7% of all our mapped supercontigs) were assigned to different chromosomes by our linkage maps and by the published chromosome map (Figure 1). These conflicting supercontig assignments were due to misassemblies that were subsequently corrected in the recently released AaegL4 genome-wide assembly (Dudchenko et al. 2017).

**Figure 1.**
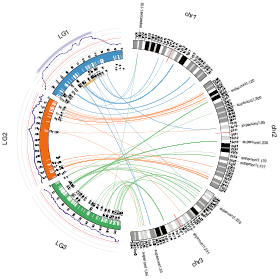
Synteny between Cross #1 linkage map and chromosome idiograms of the *Aedes aegypti* genome. Circos plot (Krzywinski, et al. 2009) shows syntenic links between linkage (left) and chromosome (right) maps. Linkage groups (LG) are 1 (blue), 2 (orange) and 3 (green). Markers are displayed with white internal ticks with position (cM) on the scale. The genetic length of linkage group 2 is over-inflated likely due to strongly distorted genotype segregation ratios in the centromeric part. Physical marker positions in Mbp refer to the AaegL4 assembly coordinates and are represented below the linkage map. The LOD curve for the sex QTL is displayed in purple in the outer track of the linkage map, with the red line representing the genome-wide statistical significance threshold. LOD of 1.5 (dark purple) and 2 (light purple) support intervals are on the top. The band 1q21 harboring the sex-determining (M) locus is represented in red. Supercontigs with conflicting locations between the genetic and the chromosome maps are shown in grey next to the chromosome map. The 63-Mbp genomic region displaying high male-female genetic differentiation in the population data is delineated with a gold strip below LG1. The 123-Mbp region with undetectable recombination between X and Y chromosomes in both intercrosses is represented by the gold and the grey strips combined.

### Intercrosses Reveal Reduced Male Recombination along a Large Region of Chromosome 1

Recombination rates were estimated by comparing the genetic distances of the linkage maps with the AaegL4 physical genomic coordinates (Supplementary Figure 1). Average recombination rate across all chromosomes was 0.90 cM/Mbp and 0.42 cM/Mbp for Cross #1 (chromosomes 1 and 3) and Cross #2 (chromosomes 2 and 3), respectively. These estimates are consistent with previously published linkage maps of the *Ae. aegypti* genome (Bonin, et al. 2015; Juneja, et al. 2014). Because sex-specific recombination rates cannot be estimated with an F_2_ intercross design, our recombination rate estimates reflect sex-averaged recombination. However, sex-specific deviations from Mendelian segregation ratios allowed us to detect sex-specific patterns of recombination in our F_2_ intercrosses.

Using a small F_2_ intercross with 22 F_2_ males and 22 F_2_ females (Cross #1), we observed significant deviations from the expected 1:2:1 segregation ratio for sections of chromosomes 1 and 2, but only chromosome 1 contained a set of markers with a sex-specific pattern of segregation (Supplementary File 2B). Namely, a 187-Mbp region of chromosome 1 spanning from 87 Mbp (37 cM) to 274 Mbp (107 cM) showed a complete lack of F_0_ paternal AA genotypes in all 22 F_2_ females, and a complete lack of F_0_ maternal BB genotypes in all 22 F_2_ males (Figure 2B-C). This deviation from Mendelian segregation ratios is expected in the SDR because markers in perfect sex linkage co-segregate with the sex-determining locus during meiosis in F_1_ males. For partially sex-linked markers, F_0_ paternal A alleles preferentially segregate in F_2_ males and F_0_ maternal B alleles preferentially segregate in F_2_ females (Supplementary File 1D). Interestingly, for one marker of this region located at position 166,482,560 bp in the AaegL4 assembly, there was a complete absence of AB heterozygotes and 42% and 58% of BB and AA homozygotes, respectively (Figure 2B). Such genotype proportions are consistent with the presence of a null allele on the Y chromosome, so that F_2_ males that inherited the F_0_ maternal B allele from their F_1_ mother were erroneously genotyped as BB homozygotes instead of AB heterozygotes. Because the probability to detect low-frequency recombinants increases with larger sample size, we further analyzed 197 F_2_ females from an independent F_2_ intercross (Cross #2). Again, we observed a complete absence of paternal F_0_ AA genotypes in all 197 F_2_ females over a 150-Mbp genomic region spanning from 61 to 211 Mbp on chromosome 1 (Figure 2D). Overall, the common region with undetectable male recombination in both Cross #1 and Cross #2 spanned 123 Mbp (from 87 Mbp to 211 Mbp).

**Figure 2.**
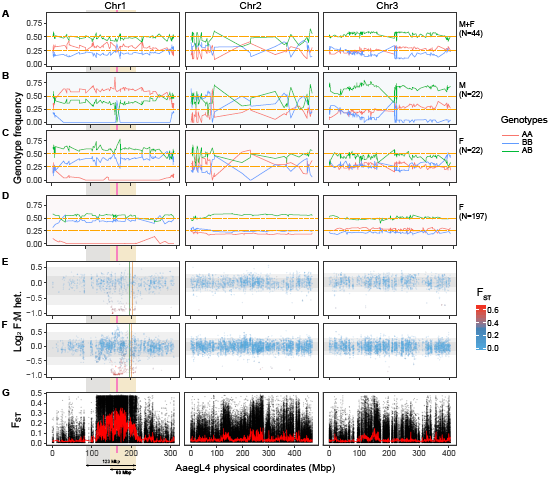
Male-female genetic differentiation and relative heterozygosity of the *Aedes aegypti* sex chromosomes. Line graphs in panels (A) through (D) represent the observed frequency of AA (red), AB (green) and BB (blue) genotypes at each marker along the 3 linkage groups. AA represents the F_0_ paternal genotype and BB represents the F_0_ maternal genotype. In Cross #1, genotype proportions of F_2_ males and females together (N = 44), males only (N = 22) and females only (N = 22) are represented in panels (A), (B) and (C), respectively. Panel (D) represents genotype proportions for 197 females in Cross #2. Scatter plots showing log_2_ female:male heterozygosity for the Brazilian and Australian mosquito populations are displayed in panels (E) and (F), respectively, with male-female *F*_ST_ values (genetic differentiation) of the corresponding markers represented in a color scale. One- and two-fold standard deviations around the mean log_2_ female:male heterozygosity are displayed on each chromosome by dark and light grey strips, respectively. Green and orange vertical lines show the genomic positions of LF284T7 and LF159T7, respectively, which are two mRNA-derived sequences mapping to cytological band 1q21 where the sex-determining (M) locus is located (Timoshevskiy et al. 2013). Differentiation values calculated for the Liverpool samples (Weir and Cockerham’s *F*_ST_) are displayed in panel (G) for each marker (dots) along each chromosome. The red line represents the average *F*_ST_ value for a 200-SNP moving window. The pink vertical line that crosses panels (B) through (F) denotes the physical position of chromosome 1 where putative null alleles were detected on the Y chromosome. The 63-Mbp genomic region displaying high male-female genetic differentiation in the population data is delineated with a gold strip below chromosome 1. The 123-Mbp region with undetectable recombination between X and Y chromosomes in both intercrosses is represented by the gold and the grey strips combined.

To confirm that the region showing reduced recombination between X and Y chromosomes contains the SDR, we employed QTL mapping in Cross #1. We found a major QTL associated with sex on chromosome 1 by standard interval mapping using a binary trait model (Figure 1). The highest logarithm of odds (LOD) score for this QTL was 7.6 at 49.7 cM with a 1.5 LOD support interval spanning from 35.8 to 114.7 cM. The genome-wide LOD threshold of statistical significance (α = 0.05) calculated from 1,000 permutation tests was 3.30. Based on the AaegL4 assembly, the genomic region associated with a significant LOD score ranged from 30.9 to 304.6 Mbp, which represents 88.7% of the chromosome 1 physical length. Markers located in this region had genotype frequencies that significantly deviated from the expected 1:2:1 Mendelian segregation ratio only when each sex was analyzed separately (Supplementary File 2B).

### X and Y Chromosomes are Genetically Differentiated Across a Large Region in the Liverpool Strain and Wild Populations

RAD markers in samples from two field-caught *Ae. aegypti* populations and WGS markers in a sample from the Liverpool strain were ordered along the three chromosomes using the AaegL4 assembly. After retaining RAD loci that were present in both sexes (< 25% missing) and polymorphic in at least one sex (minor allele frequency > 10%), the dataset from the field-caught Australian population contained 2,806, 5,103 and 4,149 SNPs on chromosomes 1, 2 and 3, respectively. A total of 329 markers were unassigned to the chromosome-wide AaegL4 assembly. The field-caught Brazilian population dataset contained 1,009, 1,909 and 1,601 SNPs on chromosomes 1, 2 and 3, respectively. A total of 140 markers were unassigned to the chromosome-wide AaegL4 assembly. In the Liverpool dataset, after retaining variants called based on a depth of coverage > 50, we analyzed 117,124 SNPs on chromosome 1, 268,602 SNPs on chromosome 2, and 179,794 SNPs on chromosome 3.

In all three *Ae. aegypti* population samples, genetic differentiation (*F*_ST_) between males and females was 3.4-to 7.1-fold higher on chromosome 1 than on the other two chromosomes (Table 1). The *F*_ST_ distributions of chromosomes 2 and 3 have > 96% overlap whereas the *F*_ST_ distributions of chromosome 1 and chromosome 2 or 3 have < 80% overlap. Genome-wide *F*_ST_ between males and females in the Liverpool strain (*F*_ST_ = 0.044) was elevated in comparison with the Australian (*F*_ST_ = 0.027) and Brazilian (*F*_ST_ = 0.026) samples. This likely reflects higher variance when estimating allele frequencies from a small pool-sequencing dataset (6-10 individuals) than from a larger individual-based dataset (> 30 individuals). Regardless of the differences in experimental protocols (pooled WGS vs. ddRADseq), sample size and geographic origin, chromosome-wide *F*_ST_ patterns were remarkably similar among all three samples (Figure 2E-G). High male-female genetic differentiation was observed across a 103-Mbp region of chromosome 1 spanning from about 111 Mbp to 214 Mbp, which represents about one third of its physical length. Six supercontigs within this region were previously mapped by FISH to bands 1p21, 1q11-14, 1q21, in close proximity to the M-locus position (1q21).

We also detected highly significant excess of heterozygosity for chromosome 1 markers in males relative to females (Figure 2E-F). Average frequency of heterozygotes (*H*) was 0.305 and 0.379 for the Australian females and males, respectively (*t-*test, *p* < 0.001), and 0.312 and 0.406 for the Brazilian females and males, respectively (*t-*test, *p* < 0.001). Differences in heterozygosity between males and females were statistically insignificant for other chromosomes in both groups, except for chromosome 3 in both the Australian sample (*t*-test, *p* = 0.030), for which females had marginally lower heterozygosity than males, and the Brazilian sample (*t*-test, *p* = 0.003), for which females had higher heterozygosity than males (Table 1). A 63-Mbp region of chromosome 1 spanning from 148 to 210 Mbp in the Brazilian population and from 150 to 211 Mbp in the Australian population contained a cluster of markers with significantly higher male-to-female heterozygosity levels. The same markers showed high male-female genetic differentiation. Interestingly, we detected a smaller region in the Australian population between 153 and 178 Mbp on chromosome 1 that contained a cluster of markers with significantly higher heterozygosity in females relative to males. This higher female-to-male heterozygosity is indicative of null alleles on the Y chromosome. This cluster of markers spans a genomic region that includes the position of a putative Y-linked null allele detected in the Cross #1 (166,482,560 bp, Figure 2B,F).

In the non-recombining XY chromosomal system, sex-linked markers should have Weir and Cockerham’s *F*_ST_ close to the maximal theoretical value of 0.5 and be homozygous in females and heterozygous in males (Brelsford, et al. 2016). Such markers were significantly more frequent on chromosome 1 when compared to the other two chromosomes (𝒳^2^ test, *p* < 0.001). Namely, markers with *F*_ST_ close to 0.5 that are homozygous in females and heterozygous in males comprised 4.4% of all markers on chromosome 1 in the Australian sample and 7.8% in the Brazilian sample. In comparison, less than 0.2% were found on chromosomes 2 and 3 for both the Australian and Brazilian samples (Table 1). Direct estimates of heterozygosity were not possible in the Liverpool sample because the pooled sequencing approach does not allow identification of individual genotypes. Instead, we considered that markers were sex-differentiated if they had *F*_ST_ > 0.4 (as in the Australian and Brazilian samples), if one allele was fixed in females (i.e., all females were homozygous) and if the allelic frequency was close to 0.5 in males. Using this approach, we detected 7.6% of sex-differentiated markers on chromosome 1, and 0.06% and 0.08% on chromosomes 2 and 3, respectively. Therefore, sex-differentiated markers in *Ae. aegypti* are robustly identified using different sequencing and analytical approaches (Table 1).

A list of markers within the differentiated XY region from the WGS experiment (Liverpool strain) would be exhaustive, but the reduced-genome-representation (ddRADseq) dataset from Australia offers insight into their location and potential effects (Supplementary File 2A). The 63-Mbp region showing high *F*_ST_ on chromosome 1 harbored 538 genes. A total of 95 unique genes located in this region were captured by 256 (16%) intragenic markers out of the total 1,616 markers assigned to this region. Of these 95 genes, 23 (24%) contained sex-differentiated markers (i.e., markers with *F*_ST_ > 0.4, heterozygous in males and fully homozygous in females). VectorBase contained significant differential expression data between males and females for 14 out of 23 (61%) of these genes at the pupal stage (Tomchaney, et al. 2014) and 11 out of 18 (61%) of these genes at the adult stage (Dissanayake, et al. 2010; Tomchaney, et al. 2014) (Supplementary File 2C). No significant enrichment of differentially expressed genes between males and females was observed on this chromosome 1 region relative to the other chromosomes. In comparison, 13 out of 23 randomly selected genes from the other two chromosomes (𝒳^2^ test, *p* = 1) were reported as differentially expressed between males and females in both studies.

DAPC performed separately for each geographic sample and chromosome showed that genetic differentiation along chromosome 1 was sufficient to assign individuals to their correct sex with 98.4% accuracy in the Australian sample and 92.4% accuracy in the Brazilian sample (Figure 3). Conversely, separation based on variation on chromosomes 2 and 3 was not better than random. Correct sex assignments were 54.8% and 45.4% for chromosome 2 and 48.4% and 51.5% for chromosome 3, in the Australian and Brazilian sample, respectively (Figure 3).

**Figure 3.**
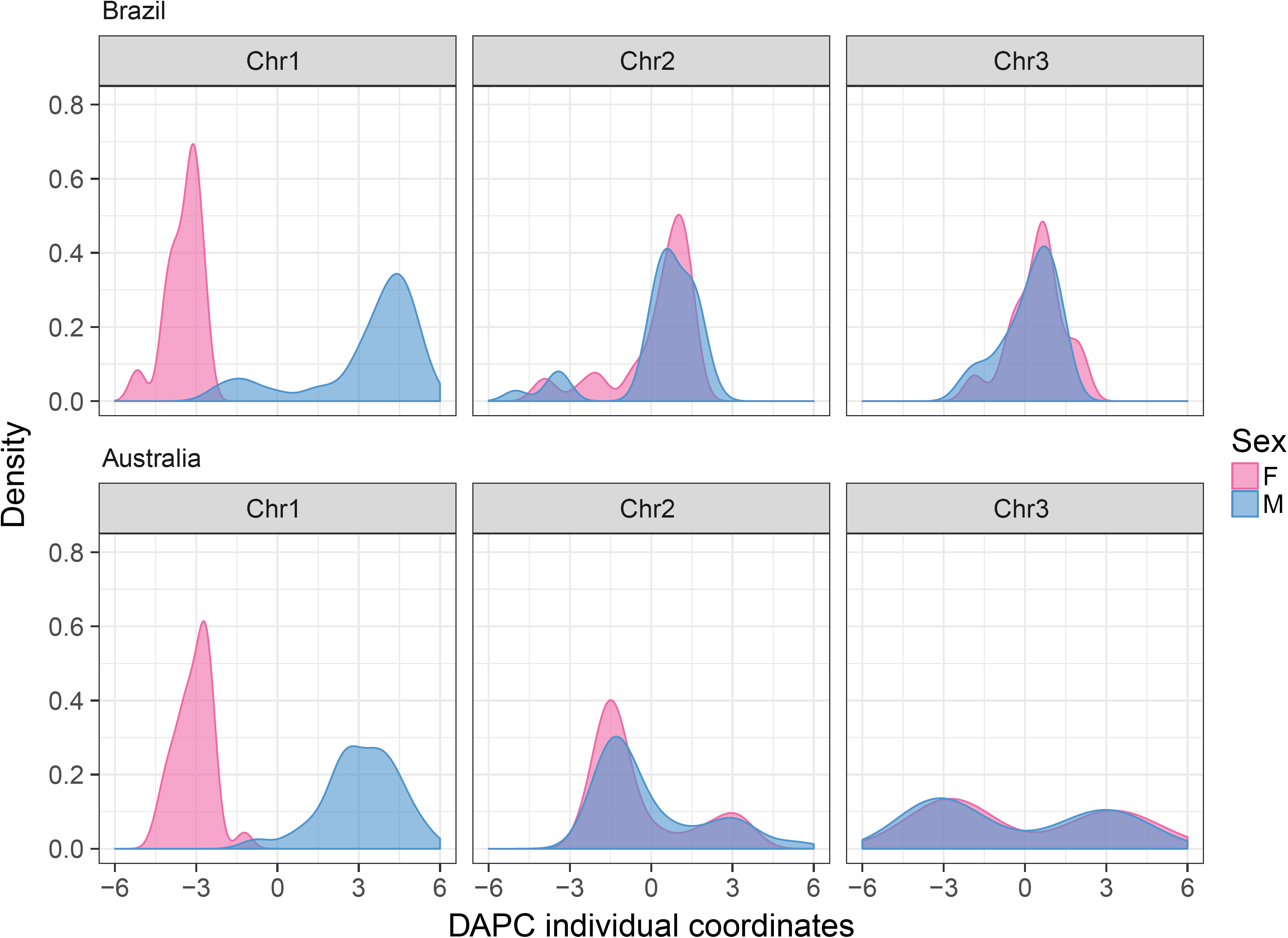
Frequency distribution of individual DAPC scores stratified by sex, for each *Aedes aegypti* chromosome and sample. DAPC accurately separates *Aedes aegypti* females from males only with chromosome 1 markers.

### Estimating Linkage Disequilibrium with the Sex-Determining Locus

Reduced recombination between X and Y chromosomes in the vicinity of the SDR is expected to lead to high LD between loci and the sex-determining locus. Because LD between such a locus (which we will denote as *A*) and the sex-determining locus (which we will denote as *M*) cannot be estimated using the standard methods for unphased autosomal genotypes, we developed an approach based on allele frequency differences between females and males.

A natural measure of LD for our purposes is 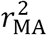, the square of the correlation between the allelic state at the sex-determining locus and the state at locus *A* in a sample consisting of equal numbers of X and Y chromosomes (e.g., sperm). The squared correlation is useful because we are not interested in the sign of the correlation (positive or negative) but only in its magnitude (large or small). Values of 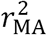 near 1 suggest that locus *A* is in the non-recombining SDR or tightly linked to it in the pseudoautosomal region. A calculation in Supplementary File 1B shows the sample value for this statistic expressed in terms of allele frequencies at locus *A* in males and females, respectively.

We used this measure of LD with the sex-determining locus for all markers across chromosome 1 and found that they show elevated LD with the sex-determining locus over about 103 Mbp in all three *Ae. aegypti* population datasets (Supplementary Figure 2G). Within this region there are small regions (tens of Mbp) that show lower levels of LD.

## Discussion

We provide compelling evidence that the sex chromosomes of the arbovirus vector, the mosquito *Ae. aegypti*, are genetically differentiated along approximately 20% of their length despite the apparent homomorphy. Our findings challenge the traditional view that the homomorphic sex-determining chromosomes in *Ae. aegypti* behave like autosomes outside a small, non-recombining SDR (referred to as the M locus in the mosquito literature). We first noticed in a small-scale F_2_ intercross (Cross #1) that recombination in male meiosis was undetectable across 40% of the chromosome 1 physical length (Figure 2A-C). Unlike previous analyses of chromosome-wide recombination rates in *Ae. aegypti* (Bonin, et al. 2015; Juneja, et al. 2014), we observed low recombination specifically in males. We next confirmed this finding in an independent F_2_ intercross (Cross #2) with a larger number of individuals (Figure 2D). Unlike a backcross design, our F_2_ intercross design did not allow a direct estimation of sex-specific recombination rates. Rather, we detected sex-specific differences in recombination patterns by measuring sex-specific deviations in Mendelian inheritance. Across the three chromosomes, a single genomic region overlapping with the sex QTL on chromosome 1 showed significant sex-specific genotype segregation bias. Genotype proportions at markers located within this region significantly deviated from the expected Mendelian segregation ratio only when samples from each sex were analyzed separately. In contrast, there was no significant difference from the expected Mendelian segregation ratio for the same markers when samples were treated regardless of sex. We observed significant deviations from Mendelian inheritance on other chromosomes but they were not sex-specific (Figure 2A-C). For instance, the centromeric part of chromosome 2 contained markers that deviated from Mendelian segregation ratios and may explain its unexpectedly large genetic length. However, these non-Mendelian segregation ratios were observed regardless of sex in the F_2_ progeny. Likewise, non-Mendelian segregation ratios were also observed on about half of chromosome 3 in both sexes. We speculate that sex-independent deviations from expected genotype segregation patterns on chromosomes 2 and 3 may have resulted from genetic incompatibilities between F_0_ parents due to the inbreeding process.

The observation of reduced male recombination across a large portion of chromosome 1 in our mapping intercrosses prompted us to examine patterns of molecular variation in natural populations reflecting more ancient recombination history. In several unrelated population samples, we found a 63-Mbp region of chromosome 1 with high male-female genetic differentiation (Figure 2E-G) and high male-to-female heterozygosity (Figure 2E-F). These features are consistent with a differentiated XY chromosomal system (Brelsford, et al. 2016), where homologous alleles are preferentially associated with either the X or the Y chromosome. The same 63-Mbp region contained two mRNA-derived sequences (LF284T7 and LF159T7) that mapped to cytological band 1q21, where the sex-determining locus is located (Timoshevskiy et al. 2013). The presence of Y-specific and X-specific alleles in a region of chromosome 1 that is in strong LD with the SDR (Supplementary Figure 2) is in line with the hypothesis that the non-recombining SDR might be expanding in *Ae. aegypti*. Furthermore, we identified a small cluster of markers inside the 63-Mbp region (between 153 and 178 Mbp) displaying high female-to-male heterozygosity consistent with null alleles on the Y chromosome (Supplementary File 2A). We did not observe a similar pattern in the Brazilian sample, but this could be due to the lower marker density. However, our interpretation is supported by one marker captured in Cross #1 that mapped to the same location and showed a genotype segregation pattern consistent with a null allele on the Y chromosome. A cluster of null alleles on the Y chromosome could be due to polymorphisms in the restriction enzyme cutting sites or to a deletion on the Y chromosome. Although we cannot distinguish between these two possibilities, both are in line with the sex chromosome evolution theory. Reduced recombination on the Y chromosome is predicted to weaken the efficiency of purifying selection and promote accumulation of deleterious mutations and subsequent degeneration of the Y chromosome. Further work, such as independent sequencing of the X and Y chromosomes, is required to test this hypothesis.

The 63-Mbp region of high male-female genetic differentiation in our population data was about two times smaller than the 123-Mbp region with undetectable male recombination in our intercrosses. In addition, there was lower male-female genetic differentiation on both sides of the 63-Mbp sex-differentiated genomic region. Outside of the 63-Mbp region, the female-to-male heterozygosity ratio was similar to that of chromosomes 2 and 3. This pattern could reflect suppression of recombination between the X and Y chromosomes that is too recent for them to have differentiated. In this case, the flanking regions would represent “evolutionary strata” analogous to those found in mammalian sex chromosomes (Lahn and Page 1999). Alternatively, the two flanking regions of the SDR may continue to recombine at a low rate, effectively preventing full differentiation between the X and Y chromosomes. Our intercrosses could have missed rare recombination events captured by the population data because of the lower number of meioses. Other studies indicate that male recombination is not completely abolished in the vicinity of the SDR (Hall, et al. 2014). Progression of homomorphic sex-determining chromosomes into heteromorphic sex chromosomes is not inevitable because examples of old homomorphic sex chromosomes exist (Abbott, et al. 2017; Charlesworth and Mank 2010; Vicoso, et al. 2013a; Yazdi and Ellegren 2014). Low but effective recombination rates can contribute to the maintenance of homomorphic sex chromosomes. For instance, extremely low but non-zero recombination rates between undifferentiated sex chromosomes in male tree frogs (*Hyla* spp.) was inferred from the population-based analyses of molecular variation, whereas laboratory crosses did not detect recombination (Guerrero, et al. 2012). Simulation work by Grossen and colleagues showed that recombination rates as low as 10^−4^ could keep sex chromosomes homomorphic (Grossen, et al. 2012).

Preservation of homomorphic sex chromosomes is generally associated with sex-biased levels of gene expression of sex-linked genes (Vicoso, et al. 2013b). Evolving sex-biased gene expression could be a mechanism that alleviates selection pressure to entirely cease recombination between chromosomal regions that contain sexually antagonistic alleles (Cheng and Kirkpatrick 2016; Vicoso, et al. 2013). This mechanism may provide another explanation for why sex chromosomes in *Ae. aegypti* are genetically differentiated under a level of recombination that is sufficient to maintain their apparent homomorphy. Perhaps mosquitoes, like birds (Vicoso, et al. 2013), have found different evolutionary solutions to deal with deleterious effects of sexually antagonistic mutations. Some lineages have maintained homomorphic sex chromosomes (*Ae. aegypti* and other Culicinae), while others evolved heteromorphic sex chromosomes (Anophelinae). Yet, the 63-Mbp region of chromosome 1 with strong male-female differentiation was not particularly enriched with genes significantly differentially expressed between males and females at either the pupal or adult stage. Further work is required to determine whether *Ae. aegypti* homomorphic sex chromosomes are nascent heteromorphic sex chromosomes or whether evolutionary mechanisms will continue to preserve their homomorphy.

It is important to note that our analyses give conservative estimates of sequence differentiation between *Ae. aegypti* sex chromosomes because any male-specific sequences without gametologs (i.e., homologous sequences on the non-recombining opposite sex chromosome) were not considered. Male-biased and male-specific sequences were identified as largely missing from the current genome assembly based on the Liverpool strain (Hall, et al. 2015; Hall, et al. 2014), and we detected such sequences in our ddRADseq datasets from wild populations (Supplementary File 1A). Long-read sequencing technology was recently used to improve the assembly of repeat-rich Y chromosome sequences in *Anopheles* mosquitoes (Hall, et al. 2016). The same approach could be used to identify additional Y-specific sequences in *Ae. aegypti* and incorporate them into the improved genome assembly. However, thousands of putative sex-differentiated markers were detected in the WGS dataset and over a hundred in the reduced genome representation (ddRADseq) dataset, demonstrating that the current *Ae. aegypti* genome sequence is still informative about the sex-specific allelic variants.

Results from our intercrosses, unrelated wild populations and the most commonly used laboratory strain all point to the commonality of genetically differentiated X and Y chromosomes in *Ae. aegypti*. This means that genetic analyses involving markers on chromosome 1 should no longer assume their pseudoautosomal behavior. Linkage mapping and genome-wide association studies should implement appropriate statistical methods for sex-linked data (e.g., XWAS (Gao, et al. 2015)). Population genetic analyses should check if marker deviations from the Hardy-Weinberg equilibrium stem from the non-autosomal nature of the chromosome 1 centromeric region. To date, population genetic analyses have proven challenging in *Ae. aegypti* as markers often show deviations the from Hardy-Weinberg equilibrium (e.g., excess homozygosity, high LD), which can be erroneously interpreted as presence of null alleles or selection signatures. Sexes should therefore always be clearly distinguished in population genetic studies and the chromosomal location of markers should be established. Where sex separation based on morphological characters is difficult (e.g., in immature stages or damaged material), DAPC with chromosome 1 markers (Figure 3) or presence of the male-specific sequences can be used (Supplementary File 1A).

Consideration of the reduced recombination along chromosome 1 in male meiosis is also warranted for vector control strategies such as the field deployment of *Wolbachia*-infected *Ae. aegypti* (Hoffmann, et al. 2015). The release stocks generally undergo several generations of backcrossing with field-derived mosquitoes to create favorable combinations of alleles that increase fitness in the field as well as in the laboratory (Hoffmann, et al. 2011). Because *Wolbachia* causes cytoplasmic incompatibility (Walker, et al. 2011), only *Wolbachia*-infected females are crossed with males from a target field population. Lower recombination in male meiosis means that males from the release colony are expected to maintain the genetic background of the field population along a significant portion of chromosome 1.

In conclusion, our discovery of a genetically differentiated homomorphic XY chromosomal system in *Ae. aegypti* lays a new foundation for the mapping and population genetic studies in this major arbovirus vector. Extensive sex-chromosome differentiation may be exploited for accurate sexing of mosquitoes with molecular markers or provide new targets for mosquito control strategies targeting the sex-determining pathway. Our finding also calls for investigation of such chromosomal features in other Culicinae mosquitoes, many of which are significant vectors of human pathogens. Thorough understanding of sex-determination mechanisms and evolution in these mosquitoes will require improved genome assemblies that should be generated separately for each sex.

## Acknowledgements

We thank Alongkot Ponlawat, Jason Richardson, three anonymous reviewers and the Lambrechts lab members for their insights. We are grateful to Eric Deveaud, Nicolas Joly, Olivia Doppelt-Azeroual and Véronique Legrand for assistance with computational analysis, and to the Nectar Research Cloud for computational resources. The opinions or assertions contained herein are the private views of the authors and are not to be construed as reflecting the official views of the United States Army, Royal Thai Army, or the United States Department of Defense. The funders had no role in study design, data collection and interpretation, or the decision to submit the work for publication.

## Supplementary Data

**Supplementary File 1. (A) Identification of male-diagnostic RAD markers**. Bioinformatic analyses undertaken to assemble male-specific scaffolds and identify male-diagnostic RAD markers are described. The resulting male-diagnostic sequences are listed in fasta format. Analytical steps are depicted in Supplementary File 1E. **(B) Method to estimate LD between a locus and the SDR. (C) Flowchart of bioinformatic analytical steps undertaken to generate a male-specific genome assembly and male-diagnostic RAD tags.** Details of the analyses are presented in the Supplementary File 1A. **(D) Diagram of the expected sex-specific segregation of marker genotypes in an F_2_ intercross.** A region of reduced male recombination depicted as a hatched area lacks AA genotypes in F_2_ females and BB genotypes in F_2_ males. The sex-determining locus is denoted as M. The A allele from the Y chromosome is shown in red font. **(E) Schematic of the experimental strategies used to build F_2_ intercross mapping populations**.

**Supplementary File 2. (A) Marker information for the population datasets.** Allelic frequencies in females (*p*_f_), males (*p*_m_) and both sexes (*p*), male-female *F*_ST_, 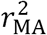, frequency of heterozygotes in females *H*_(f)_ and males *H*_(m)._ Markers are ordered according to the coordinates of the AaegL4 chromosome-wide assembly (Dudchenko et al. 2017). Relationships with the Juneja et al. (2014) linkage map (JunChr – chromosome, JunBP – base pair along a chromosome, JunSc – supercontig number, JunCM – centiMorgan position) are also represented. Each dataset is in a separate sheet. **(B) Linkage maps generated from Cross #1 and Cross #2.** Marker identification number, chromosome position, genetic position in cM, *p*-values of 𝒳^2^ tests of expected Mendelian segregation ratios, percentage of marker genotypes, marker position on supercontigs (AaegL4 assembly), and LOD scores underlying the sex QTL detection are provided. Sex-specific information is given for Cross #1. Genotype proportions were compared between males and females at each marker using 𝒳^2^ tests for Cross #1. **(C) List of intragene sex-differentiated markers in the Australian sample**. Links to the expression information curated by VectorBase.

**Supplementary Figure 1. Relationship between genetic (cM) and physical (Mbp) distances and estimated local recombination rates across the three *Ae. aegypti* chromosomes.** Physical distances in Mbp correspond to the AaegL4 chromosome-wide assembly. Chromosomes 2 and 1 were excluded from the analysis of Cross #1 and Cross #2, respectively, because of strong sex-independent deviations in genotype segregation patterns.

**Supplementary Figure 2. Estimated LD between each chromosome 1 marker and the sex-determining locus.** Individual 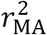 values (dots) and moving window average (line) are shown for each population sample (A: Brazil, B: Australia, C: Liverpool). Moving window averages were calculated across 20 SNPs for the Brazilian and Australian populations and 200 SNPs for the Liverpool population.

